# Enhanced olfactory memory performance in trap-design Y-mazes allows the study of novel memory phenotypes in *Drosophila*

**DOI:** 10.1101/2020.11.18.386128

**Authors:** Radhika Mohandasan, Fathima Mukthar Iqbal, Manikrao Thakare, Madhav Sridharan, Gaurav Das

## Abstract

The neural basis of behaviour is identified by systematically disrupting the activity of specific neurons and screening for loss in phenotype. Robust, high-scoring behavioural assays are thus necessary for identifying the neural circuits of novel behaviours. Here, we report the design and use of a Y-maze based classical olfactory learning and memory assay in *Drosophila*. Appetitive memory scores in our Y-mazes are considerably better and longer-lasting than that from a commonly used T-maze design. We found that the mechanism that traps flies in their choice of an odour is mainly responsible for the improving scores in the Y-mazes. Using Y-mazes, we could assay significant 24 h gustatory aversive memories in flies. These aversive memories are susceptible to protein synthesis inhibitor cycloheximide (CXM) and therefore embodies long-term memory (LTM). When anaesthesia resistant memory (ARM) deficient *radish* mutant flies are trained with dry sucrose, 24 h memory is severely disrupted. However, when we trained with 2 M sucrose-agar and tested in Y-mazes, *radish* mutants exhibited a residual 24 appetitive memory. This memory is not ARM, and we show that it is not CXM sensitive LTM either. It could be a third form of appetitive consolidated memory in flies. The Y-maze assembly described here is particularly sensitive and will thus enable the study of new memory phenotypes in *Drosophila*.

## Introduction

In flies, the link between specific neural circuit function and behaviour is established by loss or gain-of-function screens where candidate neuronal cohorts are genetically manipulated (Yoshihara and Ito, 2012; Owald, Lin and Waddell, 2015). Identifying neural mechanisms underlying behaviour becomes difficult when behaviour performance is low or erratic. Enhanced performance in behaviour assays increases the chances of observing a novel behaviour and studying its underlying neural circuitry.

Scientists have extensively studied the neural mechanisms of learning and memory in *Drosophila* (Keene and Waddell, 2007; Vosshall, 2007; Harris, 2008). Groups of flies are exposed sequentially to two sensory cues, usually odours, only one of which overlaps with a punishment or a reward. While electric shock, high-temperature or bitter taste have been employed as punishing stimuli, food, water, alcohol, or sex have acted as rewarding stimuli (Pitman *et al*., 2009; Kaun *et al*., 2011; Shohat-Ophir *et al*., 2012; Das *et al*., 2014; Galili *et al*., 2014; Lin *et al*., 2014; Masek *et al*., 2015). Subsequently, the degree of avoidance or approach elicited by the learned odour cue is quantified as a memory performance index.

In one of the earliest experiments, Seymour Benzers’ lab conditioned flies in a countercurrent apparatus to avoid odours associated with an electric shock. The same study used a Y-maze to train flies with two visible wavelengths of light (Quinn, Harris and Benzer, 1974). However, the learning scores in this pioneering work were low. Later, other groups used the countercurrent machine to train flies with sugar or shock but tested learned odour preference in a T-maze (Dudai, 1983; Tempel *et al*., 1983). Eventually, the T-maze design was modified to accommodate both training and testing of flies and memory scores improved (Tully and Quinn, 1985; Schwaerzel *et al*., 2003; Keene *et al*., 2006). A significant reason for this improvement was likely due to a shift from instrumental to classical conditioning. Flies were now unable to avoid the reward or punishment (Tully and Quinn, 1985). T-maze designs thus found wide use for assessing learning and memory in flies (Y-C Kim, Lee and Han, 2007; Colomb *et al*., 2009; Ichinose and Tanimoto, 2016). In contrast, Y-maze designs have been less commonly used for memory assays. Recently used avatars have vacuum or pressure-assisted odour streams, and they yield scores similar to that from T-mazes (Kaun *et al*., 2011; Albin *et al*., 2015; Scaplen *et al*., 2020). While effective, T-mazes can be challenging to set up and cumbersome to operate. It needs elaborate vacuum tubing arrangements, airflow, and pressure adjustments for consistent results. Getting one made requires the services of a good machine shop and can be expensive.

Here, we describe an improved setup for olfactory conditioning and testing in flies. We train flies in vials fitted with 3D printed odour cups. We test conditioned odour preference of trained flies in Y-mazes put together from fly vials, 3D printed connectors and 1 mL micropipette tips. These Y-mazes are low cost, easy to assemble, customizable and do not require vacuum or air pressure generated odour flows. They persistently yield significantly higher memory scores compared to T-mazes. Our experiments suggest that better scores arise from the trap design in Y-mazes that prevent flies from changing their preference once they enter one of the odour-choice vials.

To highlight their utility, we used Y-mazes to make a few noteworthy observations. We saw enhanced 24 h aversive memory performance with copper sulphate (CuSO_4_) reinforcement. Moreover, this memory was a protein synthesis-dependent long-term memory (LTM). The same was true for enhanced 24 h sweet taste reinforced appetitive memory too. We also observed a significant residual memory when ARM deficient *rad^MI12368-TG4.1^* mutant flies were trained with sucrose-agar and tested in Y-mazes. This residual memory was barely susceptible to CXM treatment and could be a potentially novel form of consolidated appetitive memory. Therefore, improved memory scores in the Y-mazes can foster the study of new memory phenotypes in flies.

## Materials and Methods

### Flies

All flies were reared on food containing sugar, cornflour, malt extract and yeast at 25°C and 60 % humidity, with a 12-12 h light-dark cycle in incubators. We used previously described fly strains for all experiments: Wild type Canton-S (WT-CS), *dumb^1^* and *dumb^2^* mutants (Young-Cho Kim, Lee and Han, 2007), *R58E02-GAL4 (Liu et al., 2012)*, *UAS-Shibire^ts1^* ^(Kitamoto, 2001)^ *dunce^1^* (BL6020) (Davis and Kiger, 1981), *tequila* (BL18473) (Didelot *et al*., 2006; Krashes and Waddell, 2008), *rad^MI12368-TG4.1^* (BL76737), a MiMiC-Trojan-Gal4 insertion in radish gene, creating a mutant (Lee *et al*., 2018).

### Chemicals and other materials

White oil (petroleum-derived mineral oil) of ISO VG 12 grade (Shield Lubricants), with a specific gravity of ~0.8 (23°C) was procured locally. Sucrose (ANJ biomedicals-cat no. 57-50-1), D-arabinose (MP Biomedicals-cat no.-194023), copper sulphate (CuSO_4_.5H_2_O) (SIGMA-Aldrich cat no. 209198), cycloheximide (MP Biomedicals-cat no.-100183), agar (HiMedia-cat no. GRM026), octan-3-ol (SIGMA-Aldrich-cat no. 218405), 4-methyl cyclohexanol (SIGMA-Aldrich-cat no.153095), whatman filter paper (GE-cat no. 3030-917), and polylactic acid (PLA) black 2.85 mm filament Ultimaker-cat no-1609) were procured from authorised vendors.

### Odour preparation

For Y-maze conditioning, 0.8 μl of octan-3-ol (OCT) or 0.9 μl of 4-methyl cyclohexanol (MCH) was diluted in 1.25 mL of white oil. We considered this as ~10^−3^ or 1000 fold dilution. For odour standardisation experiments, 10^−3^ dilutions were further serially diluted to ~10^−4^ or ~10^−6^. For T-maze conditioning, ~10^−3^ odour dilutions were used (8-12 μl of OCT and 9 μl MCH in 10 ml white oil).

### Olfactory associative conditioning using Y-mazes

#### a. Training setup in fly vials

5 ml of 0.75 % agar or sucrose, 50 mM to-2 M, in 0.75 % agar was poured in standard fly rearing vials (inner diameter: 23 mm, outer diameter: 25 mm, height: 90 mm) and were used for appetitive training. For aversive taste training, 80 mM CuSO_4_ +200 mM D-Arabinose in 0.75 % agar was used. For training with dry sucrose, 2.66 M sucrose solution was poured and dried on pieces of filter paper (~8.5 cm*7.3 cm). The sugar papers lined the inner side of the training vials (Supplementary Figure S2C).

20 µl odour dilutions were spotted on 1 cm * 1 cm Whatman filter paper pieces, which were placed inside custom-designed 3D printed odour cups with sieved base. The cups fit on top of the training vials (Supplementary Figure 1A, B; Figure S1A, B and supplemental video file 1). The sieve in the cup’s base allows diffusion of the odours into the tube below. We sealed the top of the odour cups with tape to prevent outward odour diffusion. Capped empty vials were used to harbour flies for 5 min between their exposure to the two odours.

#### b. Differential appetitive and aversive olfactory conditioning (training)

4-8 days old mixed-sex population of flies were used for all experiments. We starved groups of ~100 flies for ~22-24 h at 25°C in individual vials containing 0.75 % agar. We placed filter paper strips inside starvation vials to keep flies from aggregating on the agar surface and getting their wings stuck. Before training, we put ~20-30 wild-type flies that were not being used for the experiment into each training vial for 20 min to imbue the vials with ‘fly smell’. Simultaneously, the training tubes were also allowed to odorize with the appropriate odour from the odour cups.

Flies were exposed to one odour with only agar (<u>c</u>onditioned <u>s</u>timuli without unconditioned stimuli, CS^−^) and then a second odour with sugars/bitter substances in agar (CS <u>with</u> unconditioned stimuli, CS^+^). Flies were reciprocally trained such that both odours were equally used as CS^+^. Starved flies were transferred to the assigned agar vials to experience the CS^−^ odour for 5 min to initiate conditioning. The flies were then shifted to empty vials for 5 min. Next, they were flipped to the CS^+^ odour vial with either sugar-agar reward or bitter substance-agar punishment for 5 min. Any change in association times during training have been mentioned where appropriate. Food dyes could be added to the CS+ agar to monitor feeding visually. Training vials were inverted during conditioning with aversive substances and when using a shorter association time (2 min) to ensure that hungry flies find the agar bed readily due to their negative geotactic tendencies. After training, flies were maintained in 0.75 % agar vials to keep them hungry until testing after 24 h. For assaying 3, 5 or 7-day memory, flies were starved for ~6 h after training in 0.75 % agar vials. Subsequently, they were transferred to food vials until 24 h before planned testing time. They were then shifted back to 0.75 % agar vials for ~24 h of starvation until testing.

For blocking specific neuronal populations during training using the *UAS-Shibire^ts1^* transgene (Kitamoto, 2001), flies in starvation vials were kept at 32°C in a heat chamber specially designed for behavioural experiments for 30 min before training (Supplementary Figure S1E). Training was then performed at 32°C inside the chamber. After training, flies were shifted back to room temperature (22-23°C) until testing.

#### c.Y-maze testing setup

The 3D printed parts of the Y-maze were designed on the free online 3D design software TINKERCAD and printed on ULTIMAKER 2 PLUS using PLA-Black filaments. These parts include the central Y-connector with three arms, three vial connectors, and a fly loading vial (Figure 1A and S1A). The STL files for all 3D printed parts are available upon request to the corresponding author.

**Figure 1:**
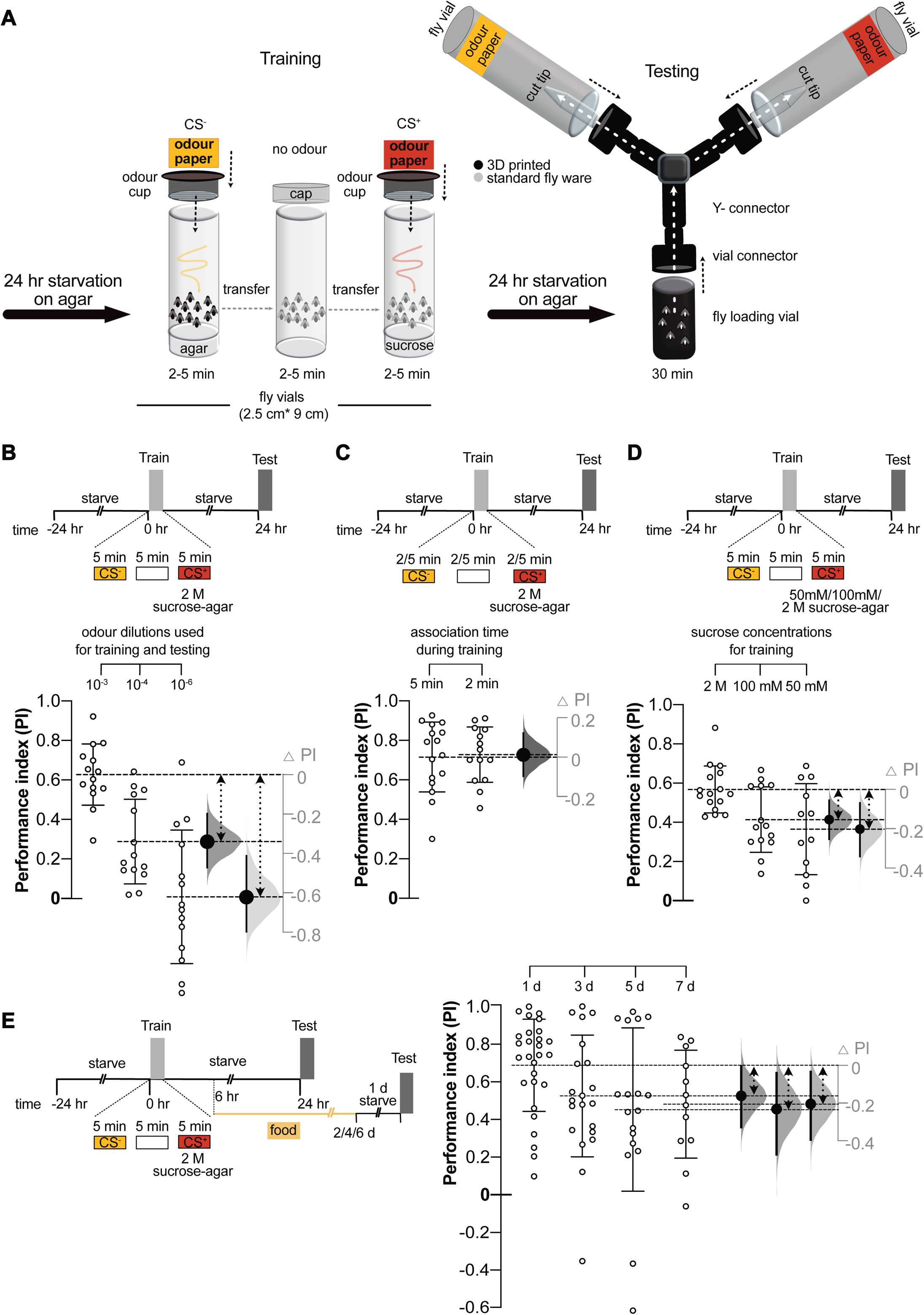
Robust and long-lasting memory performance using Y-mazes: **(A)** Classical odour conditioning of flies in vials and then testing conditioned odour preference in Y-mazes. The setups are built with readily available fly plasticware, and custom-designed 3D printed parts. **(B)** Using ~10^−3^ fold dilution of OCT and MCH for both training and testing yielded the maximum score, compared to ~10^−4^ or ~10^−6^ dilutions, *n=14; CS^+^ OCT=7, MCH=7* for all groups. **(C)** Varying odour and 2 M sucrose association time from 5 min (*n=16; CS^+^OCT=8, MCH=8*) to 2 min (*n=14; CS^+^ OCT=7, MCH=7*) during training, did not change 24 h memory performance. **(D)** Lowering sucrose concentration used for training from 2 M (*n=16; CS^+^ OCT=8, MCH=8*) to 100 mM (*n=14; CS^+^ OCT=7, MCH=7*) and 50 mM (*n=12; CS^+^OCT=6, MCH=6*) greatly reduced 24 h memory performance. **(E)** Substantial 2 M sucrose reinforced memory performance is seen even at the 7-day time point. 2 M sucrose reinforced memory performance with Y-mazes at 1 day (*n=28; CS^+^OCT=14, MCH=14*), 3 day (*n=21; CS^+^OCT=11, MCH=10*), 5 day (*n=18; CS^+^OCT=10, MCH=8*) and 7 day (*n=12; CS^+^OCT=6, MCH=6*) after conditioning. To the right of each graph, effect size or the difference in mean performance index (ΔPI) plots are shown. The curve represents the bootstrap sampling error distribution of ΔPI. The observed effect size between the experimental and a control group is depicted as a large black dot, and the 95 % CI is indicated by the ends of the vertical error bars. Double-headed dashed arrows point to the group means being compared. *Also, see supplementary Figures S1*.

The odour choice vials are the same type as the standard fly-rearing vials used for training. Whatman filter paper pieces (2.5 cm * 2 cm) was spotted with 20 µl of CS^−^ or CS^+^ odour dilutions and wedged in at the bottom of the vials. The broad end of an individual vial connector was attached to each of the two odour vials, with a 1 mL end cut tip fixed into the connector, facing inwards into the vials (Simonnet, Berthelot-Grosjean and Grosjean, 2014). The narrow end of a vial connector was fitted to individual arms of the central Y connector. The Y connector has three equal length arms that are 120 degrees apart from each other. Snug-fitting at all joints, namely those between the cut tip to vial connector, vial to vial connector, vial connectors to Y connector, is ensured by wrapping masking tape onto the surfaces (Figure S1C). The third arm of the Y connector is connected to the opaque (3D printed) fly loading vial.

The odour choice vials are wrapped in black tape to prevent the influence of non-uniform lighting on choice. They can be left uncovered if uniform lighting is ensured to both the choice arms during testing.

#### d. Conditioned odour preference of flies in Y-mazes (testing)

Before testing, we place the odour laden filter papers in the appropriate vials and assemble the Y-mazes as described. They are left to odorize for 20 min. As the strength of the volatile odours changes over time and influences choice, it is critical to carry out testing at fixed times after odours are aliquoted and Y-mazes assembled.

Previously conditioned starved flies are loaded onto the 3D printed opaque vial connected to Y-mazes. After loading, the Y-mazes are kept upright in a plastic rack (Figure S1D, E). As previously mentioned, we use a custom-designed chamber for testing (Figure S1E). The negative geotactic instinct of flies ensures that the flies climb up to the choice point through the Y connector and move towards either one of the odour choice vials. Once individual exploring flies have walked through the cut 1 ml pipette tip into one of the odour vials, they are effectively trapped and are unable to crawl back into the tip. We generally used a testing time of 30 min. In groups of flies that were tested without the tip-traps in Y-maze, flies were allowed to move freely between the two odour choice vials for the entire testing period. Testing can also be done with the Y-mazes laid horizontally. In such cases, we suggest that the choice vials are left uncovered for uniform exposure to light.

We counted the number of flies in the rest of the Y-maze assembly, other than the odour vials, to determine the proportion of flies that made a choice. We trained half of the fly groups for each condition or genotype with OCT as CS^+^ and half with MCH as CS^+^. Each group of flies, conditioned with either one of the odours, is considered ‘n=1’. Scores from any Y-maze where less than 30 flies had made an odour choice were not included in the final analysis to potentially reduce variability.

### Odour balancing in Y-mazes

A memory score is calculated by the distribution of conditioned flies between the two odour choice vials (see above). Therefore, it is critical to establish odour concentrations at which naive/unconditioned flies show an equal preference for OCT and MCH. For odour balancing, Y-mazes are prepared with concentrations of OCT and MCH that may be equally repulsive/attractive to the flies. Approximately 100 naive flies are starved for ~22-24 h on 0.75 % agar, and their choice is determined as described in the previous section. A preference index of odour choice is calculated as:

#### Preference Index = (# of flies choosing OCT*11*# of flies choosing MCH /Total number of flies that made a choice)

Hence a preference index of +1 denotes an absolute OCT preference, and −1 denotes an absolute preference for MCH. Odour concentrations are adjusted according to the observed skew to achieve an average equal preference. Odour balances vary with temperature and should be empirically determined as needed.

### Olfactory associative conditioning using T-mazes

T-maze conditioning protocols have been extensively described (Pitman *et al*., 2009; Michael J. Krashes and Waddell, 2011; M. J. Krashes and Waddell, 2011). Following is a brief description.

#### a. T-maze setup

A T-maze has three main acrylic parts. Two side walls screwed to a metal base and a middle slider block, with an elevator cavity, sandwiched in between and held with clamps (Figure S1F). It has an upper training zone and a lower testing zone that allows binary choice. A single tube connected to an odour source can be fixed at the training zone. The testing zone can take two 180° opposing tubes, each connected to either one of the odour sources. The elevator in the middle allows the transport of flies from the training to the testing zone. The training and testing zones are connected to a vacuum source, creating odour streams inside the tubes. During an experiment, air pressure is frequently checked and adjusted as needed using an airflow meter.

#### b. T-maze training and testing

Flies starved for 24 h on 0.75 % agar are trained in the upper training tube to pair CS^−^ odour with 0.75 % agar poured on a Whatman filter paper (4.8 cm * 7.5 cm) or dry filter paper for 2 min. CS^−^ odour source is then disconnected to have only room airflow through the tube for 30 seconds. Following this, the flies are transferred to the CS^+^ odour tube, lined with a filter paper with a layer of 2 M sucrose (or appropriate sugar/bitter US) in 0.75 % agar, for 2 min. Any variation in association times is noted in the manuscript.

After training, flies are transferred to 0.75 % agar for 24 h until testing. In some experiments, flies are fed briefly ~6 h after training to ensure enough flies survive until testing. For testing, flies are loaded onto the T-maze in a tube via the upper training zone and lowered to the testing zone, where they are given 2 min under uniformly illuminated conditions to choose between the opposing CS^+^ and CS^−^ odour testing tubes. The elevator is raised after 2 min, trapping the flies in the individual odour tubes. The flies are then transferred to separate vials and frozen. The number of flies choosing each odour is counted manually. A memory-driven performance index (PI) is calculated the same as with Y-mazes:

#### Performance Index = (# No. of flies choosing CS^+^ - # No. of flies choosing CS^−^)/ Total # of flies that made a choice

Historically, to negate any machine-specific odour bias, scores from two reciprocally trained fly groups tested in the same T-maze are averaged to get a single data point. In contrast, we strive to train an equal number of groups with OCT and MCH as the CS^+^ odour and consider each group of conditioned flies as n=1. Significantly, this does not alter the mean score. However, by not averaging, the standard deviation of our data sets are increased. For testing with 3D printed cone-traps (see Figure S3A), the traps are inserted into the two odour testing tubes up to a distance of 3 mm from the tube ends. Balanced odour concentrations are determined as described before commencing the experiments.

### Cycloheximide feeding

As described before, ~35 mM of cycloheximide was mixed into 0.75 % agar when flies were being starved 18 h before training (Tully *et al*., 1994; Krashes and Waddell, 2008). Cycloheximide was dissolved in warm water and subsequently mixed into melted agar before being poured and solidified in the starvation vials.

### Data plotting and Statistical analysis

Statistical analysis was performed using the software GraphPad Prism version 8.3.1. All data points are represented as a scatter plot in which each point represents a single ‘n’ with mean and ± standard deviation (SD). P values for relevant comparison of groups for all data and other relevant statistical details are reported in Supplemental Table 1 (ST1).

We plotted our data as estimation plots. These plots include a representation of the ‘effect’ size, which here is the difference in means of the compared groups (Δ <u>P</u>erformance Index-ΔPI). While the data itself is plotted using GraphPad Prism, we used the DABEST (‘data analysis with bootstrap-coupled estimation’) web application at https://www.estimationstats.com for generating the effect size plots (Ho *et al*., 2019).

The ΔPI, or the effect size axis, placed to the right of the main graphs, has its origin at the control group mean. The effect size is aligned with the experimental group mean on this axis (large black dot). The curve represents the bootstrap sampling error distribution of ΔPI. The effect size’s 95 % Confidence Interval (CI) is depicted by the two ends of the vertical error bars intersecting the effect size. The web application generates the 95 % CI and the error curve by performing 5000 bootstrap (with replacement) resamplings of the groups being compared and computing the difference in means of the resampled groups. The software also applies bias-corrected and accelerated (BCa) correction to the resampling bootstrap distributions of the effect size to account for any skew in it. Further details can be found at https://www.estimationstats.com.

## Results

### Training in vials and testing in Y-mazes can assay robust long-term memory

To simplify the handling and assembly of olfactory conditioning setup in flies, we adapted a Y-maze based design without vacuum or pressure-assisted odour streams (Figure 1A, Figure S1A-E). Please see the Materials and Methods section for details on building the Y-mazes and carrying out olfactory conditioning and odour preference testing.

We started by training flies with 2 M sucrose reinforcement and tested for 24 h memory performance, with 10^−3^, 10^−4^ and 10^−6^ dilutions of OCT and MCH, for both training and testing (Figure 1B). Naive/untrained flies prefer the two odours equally at the above concentrations (Figure S2A). We observed robust performance with 10^−3^, drastically lowered but intermediate performance with 10^−4^ and no discernible performance with 10^−6^ odour concentrations (Figure 1B). Based on our results, we decided to train and test flies with 10^−3^ dilutions of OCT and MCH for all subsequent experiments.

Flies had 30 min to choose between the two odour choice vials as this proved to be a sufficient period to allow most flies to make a choice (Figure S2B). However, we observed that a significant proportion of flies are trapped inside the choice vials by ~10-15 min (*data not shown*).

We next examined the effect of CS^+^ association time on memory performance. Decreasing 2 M sucrose with odour association time to 2 min from 5 min did not affect performance (Figure 1C). However, flies trained for 5 min with sucrose showed lower mortality upon subsequent starvation (*data not shown*). For most experiments, we subsequently preferred to use the 5 min association time due to the lower mortality rate and also because the 5-5-5 min (CS^−^-no odour-CS^+^) training regimen conveniently facilitated the conditioning of multiple groups of flies simultaneously.

We also investigated the effect of training with different concentrations of sucrose on 24 h memory scores. We observed considerably reduced memory scores when flies were conditioned with 100 mM or 50 mM sucrose compared to 2 M sucrose (Figure 1D). Since training with 2 M sucrose facilitated the highest 24 h memory performance, we used this sucrose concentration for further experiments. We also conditioned flies with dry sugar; 2.66 M saturated sucrose solution was poured and dried on filter paper (. Here, the scores were reduced compared to training with 2 M Sucrose in agar for the same association time of 5 min (Figure S2C, Figure 1C). Next, we tested the longevity of memory after a single round of training in our setup. For this, we tested 2 M sucrose-agar conditioned memory performance 1, 3, 5 and 7 days after training. We found that despite the expected decay of memory score over time, 7-day memory performance was robust (Figure 1E).

### Memory performance in our setup is dopamine signalling dependent and is also disrupted in classical memory mutants

In flies, learning and memory mutants have helped us understand the different phases of memory consolidation. The mutant *dunce* learns with sugar reinforcement but rapidly forgets (Tempel *et al*., 1983). Flies mutant for *tequila* and *rad^MI12368-TG4.1^* also show severe 24 h sucrose reinforced memory defects (Krashes and Waddell, 2008). To determine whether memory testing in Y-mazes alters dependence on specific proteins and their function, we tested the above three mutants for 24 h, 2 M sucrose reinforced memory. We found that 24 h appetitive memory was disrupted in *dunce*, *tequila* and the *rad^MI12368-TG4.1^*mutants (Figure 2A).

**Figure 2:**
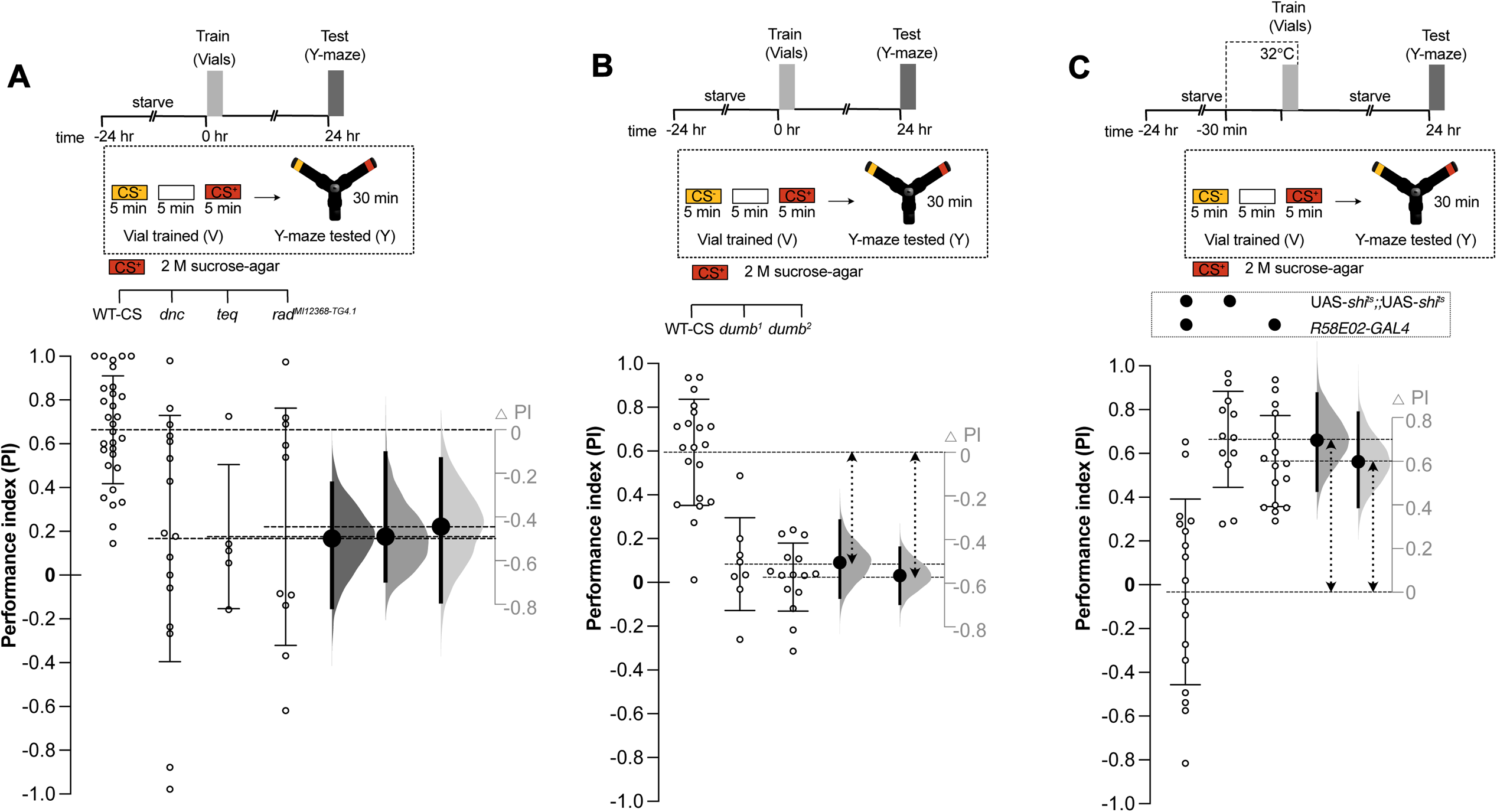
Appetitive memory formation is disrupted in mutants and is dopamine signalling dependent: **(A)** 24 h sucrose memory in *dunce (dnc)*, *tequila (teq)* and *radish (rad^MI12368-TG4.1^)* mutants. Memory is disrupted in *dnc (n=16; CS^+^ OCT=9, MCH=7), teq (n=5; CS^+^ OCT=3, MCH=5)* and *rad^MI12368-TG4.1^ (n=10; CS^+^ OCT=5, MCH=5)* as compared to WT-CS *(n=30; CS^+^ OCT=15, MCH=15)*. However, a significant residual 24 h memory is observed in *rad^MI12368-TG4.1^* mutants **(B)** 24 h memory formation is DopR1 dopamine receptor-dependent. Compared to WT-CS flies *(n=20; CS^+^ OCT=10, MCH=10)*, appetitive memory performance is abolished in *dumb^1^* (*n=8; CS^+^ OCT=5, MCH=3*) and *dumb^2^* (*n=15; CS^+^ OCT=7, MCH=8*) mutants. **(C)** Activity of the dopaminergic PAM cluster is required for appetitive memory formation in vials. Flies were trained with 2 M sucrose reinforcement at 32°C to block PAM dopaminergic neurons during training. *R58E02-GAL4* drove *UAS-Shibire^ts1^* transgene in reward PAM dopaminergic neurons *(n=17; CS^+^ OCT=9, MCH=8*). Memory performance was severely impaired in these flies compared to the two parental lines, *UAS-Shibire^ts1^(n=12; CS^+^ OCT=6, CS^+^ MCH=6)* and *R58E02-GAL4 (n=16;CS^+^ OCT=8, CS^+^ MCH=8)*. Effect size or the difference in mean performance index (ΔPI) plots are shown for all. They show the observed effect size of the experimental group compared to a control (large black dots), 95 % CI (end of vertical error bars) and the bootstrap resampling error (curve) of effect size. Double-headed dashed arrows mark the groups being compared.

Formation of appetitive memory requires dopamine release from the neurons of the PAM (protocerebral anterior medial) cluster and functional DopR1 dopaminergic receptor in the mushroom body (Young-Cho Kim, Lee and Han, 2007; Burke *et al*., 2012; Liu *et al*., 2012). To test the dopamine dependence of memory formed in our setup, we first tested 2 M sucrose reinforced 24 h memory in two null mutant strains for DopR1, namely *Dumb^1^* and *Dumb^2^* (Young-Cho Kim, Lee and Han, 2007). Olfactory memory performance was abolished in the *Dumb^1^* and *Dumb^2^* mutants compared to control (Figure 2B).

Next, to test the role of the PAM dopaminergic neurons directly, we expressed the dominant temperature-sensitive *UAS-Shibire^ts1^* transgene (Kitamoto, 2001) in almost all PAM neurons using the *R58E02-GAL4* driver line (Liu *et al*., 2012; Huetteroth *et al*., 2015; Yamagata *et al*., 2015). The *UAS-Shibire^ts1^* transgene allowed us to transiently silence the rewarding PAM dopaminergic neurons at the restrictive temperature of 32°C during training with 2 M sucrose. Upon testing at 24 h, we saw that the experimental flies expressing *UAS-Shibire^ts1^* in *R58E02-GAL4* labelled neurons exhibited no apparent memory performance (Figure 2C). Notably, a memory defect was not observed in the identical genotype when the experiment was carried out at the permissive temperature of 24°C (Figure 2D). The above results ascertained that learning in our assay setup required dopamine release from the PAM neurons and functional DopR1 receptor expressed in the mushroom body neurons.

### Testing in the Y-mazes is key to improved scores over the T-maze

We noticed that 24 h memory performance in Y-mazes (mean PI=~0.6-0.8) were considerably higher than reported using T-mazes (Tully *et al*., 1994; Michael J. Krashes and Waddell, 2011; M. J. Krashes and Waddell, 2011). To carry out a side by side comparison with a T-maze based protocol, we trained groups of flies with 1 M sucrose (in agar) reinforcement in vials or T-maze tubes for the same CS^+^ association time of 2 min. From both training conditions, half of the groups were tested in Y-mazes, and the rest in T-mazes for 24 h memory. Hence, we had four groups in each experiment; a) vial-trained and Y-maze tested (VY), b) vial-trained and T-maze tested (VT), c) T-maze tube trained and Y-maze tested (TY) and d) T-maze tube trained and T-maze tested (TT) (Figure 3A, schema).

**Figure 3:**
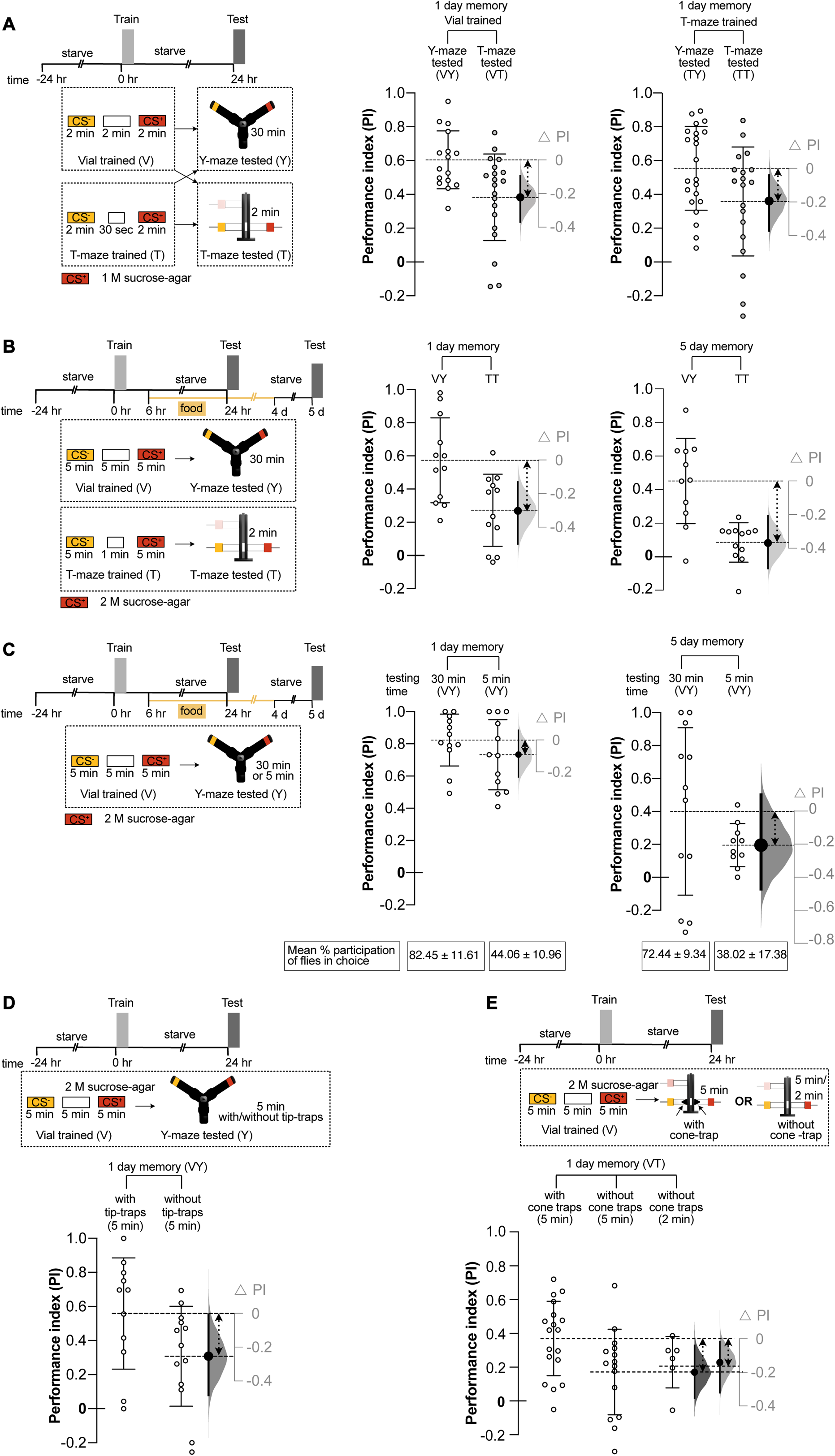
Olfactory memory assay using Y-mazes markedly improves scores over T-maze based assaying. **(A)** Irrespective of training condition, in vials or in T-maze tubes, testing in Y-mazes yields superior scores to that of the T-mazes. The Vial trained and T-maze tested group *(VT; n=20; CS^+^ OCT=11, MCH=9)* has considerably reduced performance from the Vial trained and Y-maze tested group (*VY; n=16; CS^+^ OCT=8, MCH=8)*. Similarly, the T-maze tube trained and T-maze tested group (TT; *n=19; CS^+^ OCT=9, MCH=10*) also has reduced performance from the T-maze tube trained and Y-maze tested group *(TY; n=22; CS^+^ OCT=11, MCH=11)*. **(B)** Significant 1 day scores are seen in both the Vial trained and Y-maze tested group (VY) and the T-maze tube trained and T-maze tested group (TT). However, the VY group *(1 day; n=12; CS^+^ OCT=6, MCH=6)* has substantially higher memory performance than the TT group *(1 day; n=12; CS^+^ OCT=6, MCH=6)*. A similar trend manifests on day 5 for the two groups. However, while VY (5 day; *n=11; CS^+^ OCT=5, MCH=6*) memory performance is still robust, TT group *(5 day; n=12; CS^+^ OCT=6, MCH=6)* memory performance is close to zero. **(C)** 1 day after training, testing for 5 min *(VY, 5 min; n=12; CS^+^ OCT=6, MCH=6)* has very little reduction in memory performance compared to the 30 min (VY. 30 min; *n=12; CS^+^ OCT=6, MCH=6)* tested group in Y-mazes. 5 days after training, a modest but larger reduction was seen in the 5 min tested group *(VY, 5 min; n=10; CS^+^ OCT=4, MCH=6)* compared to 30 min tested group *(VY, 30 min; n= 12; CS^+^ OCT=6, MCH=6)*. **(D)** 1 day memory performance of flies trained with 2 M sucrose-agar in vials but tested for 5 min in Y-maze with/without tip-traps. Reduced 1 day memory performance is observed in flies tested in Y-maze without tip-traps *(VY, 5 min; n=13; CS^+^ OCT=6, MCH=7)* when compared to flies tested in Y-maze with tip-traps *(VY, 5 min; n=11; CS^+^ OCT=5, MCH=6)*. **(E)** 1 day memory performance of flies trained with 2 M sucrose-agar in vials but tested in T-maze with/without 3D printed cone-traps. 1 day memory performance improved in flies tested in T-maze with cone-traps for 5 min *(VT, 5 min; n=18 CS^+^ OCT=10, MCH=8)* when compared to flies tested in T-maze without cone-traps for 5 min *(VT, 5 min; n=15; CS^+^ OCT=8, MCH=7)* or 2 min *(VT, 2 min; n=6; CS^+^ OCT=3, MCH=3)*. Effect size or the difference in mean performance index (ΔPI) plots are shown for all. They show the observed effect size of the experimental group compared to a control (large black dots), 95 % CI (end of vertical error bars) and the bootstrap resampling error (curve) of effect size. Double-headed dashed arrows mark the compared groups.

Regardless of the training condition, we observed that the groups tested in the Y-mazes showed consistently higher mean performance than those tested in the T-mazes (VY > VT and TY>TT). This result strongly suggested that testing in the Y-maze assembly was responsible for the higher scores obtained. Also, groups tested in the same apparatus had similar mean scores regardless of the training conditions (VY≈TY and VT≈TT), again highlighting the importance of testing paradigms in the magnitude of memory performance (Figure 3A).

Next, we directly compared the longevity of memory performance using Y-mazes and T-mazes. Fly groups were trained in vials and T-maze tubes with 2 M sucrose for 5 min. The memory of both these groups was tested one day and five days after training with the vial trained flies tested in Y-mazes (VY) and the T-maze tube trained flies tested in T-mazes (TT).

We found that, as before (Figure 1D), significant 1-day and 5-day memory performance was seen in VY groups. In comparison, TT groups showed considerably reduced memory scores at both time points (Figure 3B). This outcome suggested that workable memory scores were hard to get with the T-mazes beyond five days, whereas robust 7-day memory performance was attainable with Y-mazes (Figure 1D). Therefore, the Y-maze assemblies yielded superior memory scores over a longer time too.

### Trapping flies improved scores in the Y-mazes

Next, we considered the possible parameters that could aid in improving memory retrieval in the Y-mazes. Firstly, we wondered whether increasing the memory testing time from 2 min in T-maze to 30 min in Y-maze could result in higher memory scores. To test this, we assayed the memory performance of flies in the Y-maze for 30 min and 5 min both one day and five days after training with 2 M sucrose (Figure 3C, schema).

Compared to 30 min testing, mean memory scores upon 5 min testing was slightly reduced after one day, and the trend became pronounced after five days (Figure 3C). We also calculated the mean per cent of flies that made a choice (trapped in odour vials) in 30 versus 5 min testing. We observed that participation was significantly reduced for 5 min testing compared to 30 min testing, one day and five days after training (see mean participation values in Figure 3C). Thus increasing testing time may result in better performance, especially when testing long-term memory beyond 2-3 days. However, even with a short 5 min testing, performance in the Y-mazes was consistently higher than in T-mazes under similar conditions (compare 5 min VY groups in Figure 3C with TT groups in Figure 3B).

Secondly, we tested the role of the end-cut pipette tips that trap flies inside the odour vials. For this, we assayed 2 M sucrose reinforced memory performance one day after training, with and without the tip-traps. We restricted the testing time to 5 min to keep the results comparable with T-maze testing times (Schema Figure 3D). Here, we found that testing without the tip-traps significantly reduced memory performance (Figure 3D). This result suggested that the tip-trap design plays a major role in obtaining superior memory scores in the Y-mazes. When we repeated this experiment, now with 30 min testing, there was a complete disruption of memory performance (Supplementary Figure S3A). To further investigate the contribution of trapping flies, we introduced a similar trap design into the T-mazes. We used 3D printed conical traps that fit into the testing arms of the T-mazes. These cones have a small opening at the narrow end, mimicking the ‘tip-traps’ used in the Y mazes (Supplementary Figure S3B). We trained flies with 2 M sucrose in vials (schema, Figure 3E) and tested memory performance after 24 h in T-mazes, with and without the conical traps. We found that 5 min testing with the traps considerably improves memory scores over the control group tested without traps.

Similar improvement was seen compared to groups tested for 2 min without traps (Figure 3E). We did not carry out 2 min testing with tip-traps as not enough flies committed to a choice in this period. Overall, these results suggest that during odour memory testing, trapping the flies once they have committed to a choice improves memory scores.

### 24 h sweet and aversive taste memories are protein synthesis-dependent

Previously, it had been difficult to see any bitter or aversive taste reinforced memory scores 24 h post-training in flies (Das *et al*., 2014). We reasoned that T-maze conditioning protocols that do not typically yield scores at a particular time point might do so with Y-maze testing. We, therefore, verified whether assaying aversive taste learning in Y-mazes could enable the study of long-term taste aversion.

Copper sulphate (CuSO_4_), a compound listed as a pesticide, is toxic to flies (Živanov-Čurlis *et al*., 2006; Alaraby, Hernández and Marcos, 2017; Balinski and Woodruff, 2017; Halmenschelager and da Rocha, 2019). Our unpublished data shows that flies avoid it. To determine whether 24 h aversive memory can be detected in Y-mazes, we conditioned flies in vials with 80 mM CuSO_4_ mixed with 200 mM D-Arabinose, as a carrier to promote ingestion of CuSO_4_. D-Arabinose is a sweet but non-metabolisable sugar (Burke and Waddell, 2011). Control flies were parallelly conditioned with only 200 mM D-Arabinose. These flies were tested 24 h later in Y-mazes (VY groups). To directly compare with T-maze conditioning, identically treated flies were trained and tested in T-mazes (TT groups).

We observed that training and testing in the T-maze yielded a low positive score with 200 mM D-Arabinose and a low negative score with 80 mM CuSO_4_ + 200 mM D-Arabinose (Figure 4A). When mixed appetitive and aversive stimuli are used for training, the memory score is the sum of the parallel and opposing memories reinforced by the two components (Das *et al*., 2014). Therefore, the difference between the means of the two groups (D-Arabinose versus CuSO_4_+ D-Arabinose) measures CuSO_4_ reinforced aversive memory. This difference is modest in the TT groups and makes it difficult to investigate the aversive memory further (Figure 4A). In contrast, the approach score with 200 mM D-Arabinose and the avoidance score with 80 mM CuSO_4_ + 200 mM D-Arabinose are greater in the VY groups. The difference between their means is also substantial (Figure 4A). We concluded that Y-maze testing could enhance aversive memory scores, making it possible to study the underlying neural mechanisms.

**Figure 4:**
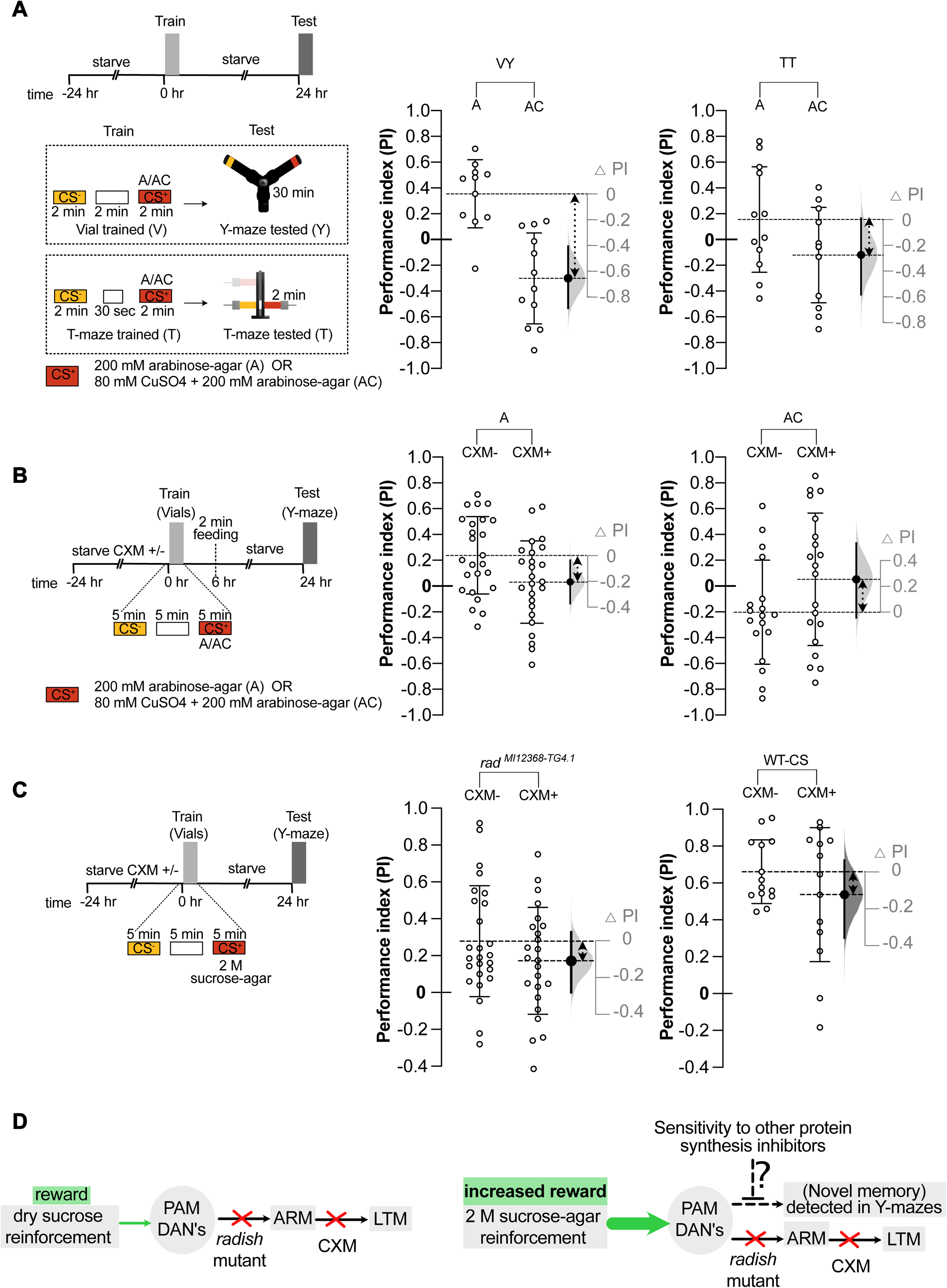
Y-mazes enable studying of novel memory phenotypes: **(A)** Enhanced CuSO_4_ reinforced aversive memory is seen in the Y-maze tested groups (VY) compared to the T-maze tested group (TT). In the Y-maze tested groups, the difference in memory performance between 200 mM D-Arabinose training *(VY-A; n=12; CS^+^ OCT=6, MCH=6)* and the 80 mM CuSO_4_ + 200 mM D-Arabinose training (*VY-AC; n=12; CS^+^ OCT=6, MCH=6)* was substantial. In contrast when the same experiment was done with T-mazes, memory performance difference between 200 mM D-Arabinose training *(TT-A; n=11; CS^+^ OCT=5, MCH=6)* and 80 mM CuSO_4_ + 200 mM D-Arabinose training *(TT-AC; n=12; CS^+^ OCT=6, MCH=6)* was modest. **(B)** To ascertain protein synthesis dependence of robust aversive taste memories, flies were trained in vials and tested for 24 h memory performance in Y-mazes. 200 mM D-Arabinose reinforced appetitive memory performance is reduced upon cycloheximide treatment *(A-CXM^+^; n=25; CS^+^ OCT=12, MCH=13)* compared to the control *(A-CXM^−^; n=25; CS^+^ OCT=12, MCH=13)*. Further, 80 mM CuSO_4_ + 200 mM D-Arabinose reinforced aversive memory performance upon cycloheximide treatment *(AC-CXM^+^; n=18; CS^+^ OCT=9, MCH=9)* was also reduced compared to the control *(AC-CXM^−^; n=20; CS^+^ OCT=10, MCH=10)*. To increase survival, all flies were briefly fed for 2 min on 2 M sucrose, 6 h after training. **(C)** To ascertain whether 24 h appetitive memory upon 2 M sucrose reinforcement is protein synthesis dependent LTM or anaesthesia resistant memory, *rad^MI12368-TG4.1^* mutants and WT-CS were treated with CXM. Reduction in memory is observed in *rad^MI12368-TG4.1^* CXM^+^ groups (*n=24; CS^+^ OCT=11, MCH=13*) compared to *rad^MI12368-TG4.1^* CXM^−^ groups (*n=26; CS^+^ OCT=13, MCH=13*). However, CXM resistant memory (mean PI=~0.18) remained in the *rad^MI12368-TG4.1^* CXM^+^ groups. Reduction in memory is also observed in WT-CS CXM^+^ groups (*n=13; CS^+^ OCT=7, MCH=6*) compared to WT-CS CXM^1^groups (*n=14; CS^+^ OCT=7, MCH=7*). **(D)** Model explaining different memory traces, consolidated upon dry sucrose and sucrose-agar reinforcement. To the right of each graph, effect size or the difference in mean performance index (ΔPI) plots are shown. These plots depict the bootstrap sampling error distribution curve of ΔPI. The actual mean difference of the groups is depicted as a large black dot and the 95 % CI is indicated by the ends of the vertical error bars. Double-headed dashed arrows between the group-means indicate the groups being compared.

Because we could see a robust CuSO_4_ reinforced 24 h aversive memory using Y-mazes, we investigated whether such memory is protein synthesis-dependent. We placed experimental flies on 35 mM of cycloheximide (CXM), a protein synthesis inhibitor, mixed in 0.75 % agar during the ~17 h starvation period before training. As before, we conditioned flies with 200 mM D-Arabinose and 80 mM CuSO_4_ + 200 mM D-Arabinose and then tested for 24 h memory. They were briefly fed for 2 min with 2 M sucrose, 6 h after training, to keep enough flies alive in the wake of cycloheximide exposure. We observed that both 200 mM D-Arabinose memory and 80 mM CuSO_4_ + 200 mM D-Arabinose reinforced memories were reduced in the CXM fed (CXM^+^) group compared to the control (CXM^−^) group. We concluded that both D-Arabinose reinforced appetitive and CuSO_4_ reinforced aversive memories were sensitive to CXM treatment and thus likely to be protein synthesis dependent (Figure 4B).

### Presence of a 24 h appetitive memory in flies which is neither ARM nor LTM

We observed a 24 h residual memory in *rad^MI12368-TG4.1^* mutant flies upon 2 M sucrose-agar training (Figure 2A). However, in agreement with earlier studies (Krashes and Waddell, 2008), we saw no such residual memory upon training *rad^MI12368-TG4.1^* mutant flies with dry sucrose and testing in the Y-mazes (Figure S4A). Therefore, we investigated the protein synthesis dependence of this residual 24 h appetitive memory in *rad^MI12368-TG4.1^* flies. Mutant *rad^MI12368-TG4.1^* flies were fed CXM as mentioned above, following which they were trained with 2 M sucrose-agar. Upon testing in Y-mazes, we found a mild disruption of 24 h appetitive memory in CXM^+^ groups compared to control CXM^−^groups, in the mutant flies (Figure 4C). The still leftover memory trace in *rad^MI12368-TG4.1^* mutants is neither anaesthesia resistant memory (ARM) nor protein synthesis dependent LTM which is disrupted upon CXM treatment. Control WT-CS flies trained with 2 M sucrose-agar also showed a mild disruption in memory upon CXM treatment (Figure 4C). However, upon dry sucrose training, CXM mediated disruption of memory was much more severe in the Y-mazes (Figure S4B), similar to what had been reported before (Krashes and Waddell, 2008). Our model is outlined in Figure 4D.

## Discussion

### Y-mazes for olfactory behaviours

The development of reliable learning and memory assays enabled the neurogenetic study of fly learning and memory (Quinn, Harris and Benzer, 1974; Tully *et al*., 1994; Pitman *et al*., 2009; Tabone Christopher and de Belle, 2014). The Y-mazes we have described here have the potential to uncover novel memory phenotypes because of their superior performance. They are easy to build and cost a fraction of other conditioning setups currently in use. The lack of vacuum-assisted odour streams does away with tubings, clamps, vacuum pumps, and airflow monitoring, making handling easy.

Miniature Y-mazes have been used in flies for measuring working memory and exploratory behaviour (Buchanan, Kain and de Bivort, 2015; Lewis *et al*., 2017; Cleal *et al*., 2020). In one of the earliest conditioning experiments, flies were trained in one arm of a Y-maze to avoid a specific wavelength of light paired with aversive quinine taste. However, scores in the visual cue reinforced assay was low (Quinn, Harris and Benzer, 1974). Y-mazes without trap design and active competing airflows have also been used recently for testing learned odour preference of flies previously conditioned with sugar or alcohol (Kaun *et al*., 2011; Albin *et al*., 2015; Nunez, Azanchi and Kaun, 2018). However, memory scores from the above Y-mazes are lower than the scores we report here. Y-mazes and T-mazes have also been used to measure olfactory choice, acuity and habituation in flies (Rodrigues and Siddiqi, 1978; Chakraborty, Goswami and Siddiqi, 2009; Das *et al*., 2011; Simoes, Ott and Niven, 2011; Simonnet, Berthelot-Grosjean and Grosjean, 2014).

### Y-mazes against T-mazes

We have compared our Y-mazes directly to T-mazes (Tully and Quinn, 1985). We found that memory scores in Y-mazes were markedly higher than in T-mazes (Figure 3, 4).

We also observed durable 7-day appetitive memory scores in the Y-mazes (Figure 1E). In contrast, T-maze memory scores were already lower than 0.1 after five days (Figure 3). A similar drop in memory scores by day five have been seen in T-mazes (Krashes and Waddell, 2008; Colomb *et al*., 2009). Long-lasting 7-day memory scores in T-mazes have only been reported upon artificial induction of appetitive memory (Huetteroth *et al*., 2015).

Our experiments suggest that testing in the Y-mazes is key to improved scores. We tested the role of the duration of testing and the trapping of flies, in enhancing scores. Our results showed that decreasing choice time from 30 min to 5 min in the Y-maze diminished both the number of flies trapped in the choice vials and overall scores (Figure 3C). Nevertheless, this is unlikely the primary reason for improved scores, as 5-min scores in the Y-maze, is still greater than comparable T-maze scores.

Our investigations indicate that the trap design that prevents change in preference once a choice has been made (Simonnet, Berthelot-Grosjean and Grosjean, 2014), has a major role in improving scores. Testing flies without tip-traps drastically reduced Y-maze memory scores (Figure 3D, Figure S3A). Further, memory scores in T-mazes were also improved when a similar trapping mechanism was introduced (Figure 3D). Other factors like reduced crowding at the choice point (Quinn, Harris and Benzer, 1974) or lack of airflow in the setup could also have minor roles in improving scores.

### The Y-mazes are versatile

Testing in Y-mazes can be coupled to other training protocols. For example, electric shock conditioned flies can be tested in Y-mazes to improve test scores. We show that Y-mazes are compatible with the thermogenetic silencing of neurons. It is easy to carry out both training and testing at 32°C by placing the training vials and the Y-mazes inside a temperature control chamber (Figure S1F). As flies are trained in transparent vials, optogenetic manipulations during training are possible. The testing setup can be made optogenetics-ready by using transparent Y-connectors. As our Y-mazes do not require a vacuum source, they are portable and can be used for science outreach outside the laboratory.

### Revealing novel memory phenotypes with the Y-mazes

Long-term memory (LTM) in animals including fruit flies, is operationally defined by its susceptibility to drugs that disrupt protein synthesis (Davis and Squire, 1984; Hernandez and Abel, 2008). Flies trained with a single round of shock exhibit sufficient 3 h memory but weak 24 h memory (Quinn, Harris and Benzer, 1974; Tully and Quinn, 1985; Tully *et al*., 1994). Such a memory does not seem to have an LTM component as it was not sensitive to the feeding of the protein synthesis inhibitor cycloheximide (CXM) (Tully *et al*., 1994). Significant 24 h aversive LTM memory formation requires multiple rounds of spaced training in fed flies (Tully *et al*., 1994). Interestingly, two distinct additive memory traces are formed after spaced training; a CS^+^ linked aversive memory and a CS^−^ linked appetitive ‘safety’ memory. Only the CS^−^ linked 24 h safety memory is CXM sensitive and thus represents the LTM component (Jacob and Waddell, 2020).

Single round shock training can also give rise to aversive 24 h LTM when flies are trained hungry and then fed after training (Hirano *et al*., 2013). In contrast, if hungry flies are kept starving after spaced shock training, LTM memory is impaired and only CXM insensitive memory is formed (Plaçais and Preat, 2013). It has thus been suggested that hunger biases the brain away from aversive LTM formation (Plaçais and Preat, 2013).

When flies undergo a single round of odour-conditioning with bitter-tasting compounds, the resulting memories decay rapidly (Keene and Masek, 2012; Das *et al*., 2014). Such rapid decay dynamics made it difficult to ascertain protein-synthesis dependence for bitter compound reinforced memories. In a recent study (Chakraborty *et al*., 2021) and in this work, we report the formation of durable bitter compound reinforced 24 h aversive memory in flies. Here, we found that 24 h aversive memory reinforced by 80 mM CuSO_4_ (with 200 mM D-Arabinose carrier) in hungry flies was severely impaired upon CXM treatment. A similar phenomenon was seen in the case of conditioning with 200 mM D-Arabinose alone (Figure 4A and 4B). Thus, we made two important observations. First, in contrast to an earlier study, aversive LTM could form even when flies are starved before and after training (Plaçais and Preat, 2013). One caveat is that we briefly fed the flies for 2 min with 2 M sucrose, 6 h after training. Second, that sweet taste alone may reinforce appetitive 24 h LTM. This was not possible to test earlier because robust D-Arabinose reinforced memory was not reported 24 h after training.

When wild type flies are trained with dry sucrose, the resultant 24 h memory is likely to be mostly the nutrient-value reinforced component (Das *et al*., 2014; Huetteroth *et al*., 2015). This same 24 h memory was almost completely abolished upon CXM treatment in both T-mazes (Krashes and Waddell, 2008) and Y-mazes (Figure S4B). However, when training was done with sucrose-agar and tested in Y-mazes, we find that CXM treatment shows only a small decrease in memory performance (Figure 4C). This could be because flies consume a lot more from sucrose-agar than dry sucrose during training and the resulting increase in reward may reinforce multiple memory components that exist parallelly (model, Figure 4D). Indeed, even training with non-nutritious sugars in agar can give rise to robust 24 h memory (Figure 4A and 4B, *data not shown*).

Previous studies have shown that, in flies trained with dry sucrose, LTM formation is dependent on ARM consolidation. Hence, *radish* mutants trained with dry sucrose show no discernible 24 h memory in both T-mazes (Krashes and Waddell, 2008) and Y-mazes ( Figure S4A). However, in *rad^MI12368-TG4.1^* mutant flies trained with 2 M sucrose-agar exhibits a residual 24 h memory when tested in Y-mazes (Figure 2A). This residual memory in *rad^MI12368-TG4.1^* mutant flies was slightly sensitive to protein synthesis inhibition by CXM (Figure 4C). In agreement with the existing model, this shows that LTM memory is mostly disrupted in radish mutant flies. We speculate that the CXM resistant residual memory in *rad^MI12368-TG4.1^* mutant flies could represent a third kind of memory trace that might be susceptible to other protein-synthesis inhibiting compounds (model, Figure 4D). Indeed, in honeybees, more than one kind of LTM has been defined based on their vulnerability to different protein-synthesis inhibiting compounds (Eisenhardt, 2014). As discussed before, the increased reward value from sugar-agar compared to dry sugar could be responsible for forming multiple memory components with different kinetics and drug sensitivities.

In conclusion, we have highlighted the benefits of a newly designed Y-maze assembly, the most important of which is improved memory scores. These Y-mazes will enable us to study novel memory phenotypes more easily. Potential users can also easily modify parts of the assembly to suit their needs. We anticipate that the Y-maze assembly described here will be widely adopted by researchers investigating mechanisms of learning and memory in flies.

## Acknowledgements

We would like to thank all our colleagues in the brain and feeding behaviour laboratory, NCCS, for their support, help with experiments and feedback on the manuscript. We also thank Suewei Lin, Vincent Croset, Emmanuel Perisse, Johannes Felsenberg, Wolf Huetteroth and Scott Waddell for their critical reading of the manuscript and for providing valuable feedback and insights. We thank Paola Cognini our vial training paradigm has been inspired by her similar design. We thank Rajkumar Pawar for handling the lab purchases, for making fly food and for stock maintenance. We thank Umesh Chachala and Abhay Jadhav for washing fly bottles and vials. RM is funded by a PhD fellowship from NCCS. MRT is funded by a Department of Biotechnology (DBT) doctoral scholarship. FMI and MS were hired from a Ramanujan Fellowship to GD. GD is a Ramanujan Fellow (SB/S2/RJN-048/2017) awarded by the Science and Engineering Research Board (SERB), part of the Department of Science & Technology (DST), Government of India. The lab is also funded by a Core Research Grant (CRG/2019/005587) from SERB and generous intramural funding support from NCCS.

## Contributions

G.D. and R.M. conceived the design of this particular Y-maze. Initial trials and design refinement was done by R.M., M.S. and G.D. All experiments described here were performed by R.M., F.M.I and M.T. G.D. directed the research and wrote the manuscript with inputs from R.M.

## Data availability

All raw data, including STL files for the Y-maze parts, are available upon request to the corresponding author.

