## Supplementary Material for "Enhanced olfactory memory performance in trap-design Y-mazes allows the study of novel memory phenotypes in *Drosophila*"

**Supplementary Figure 1**

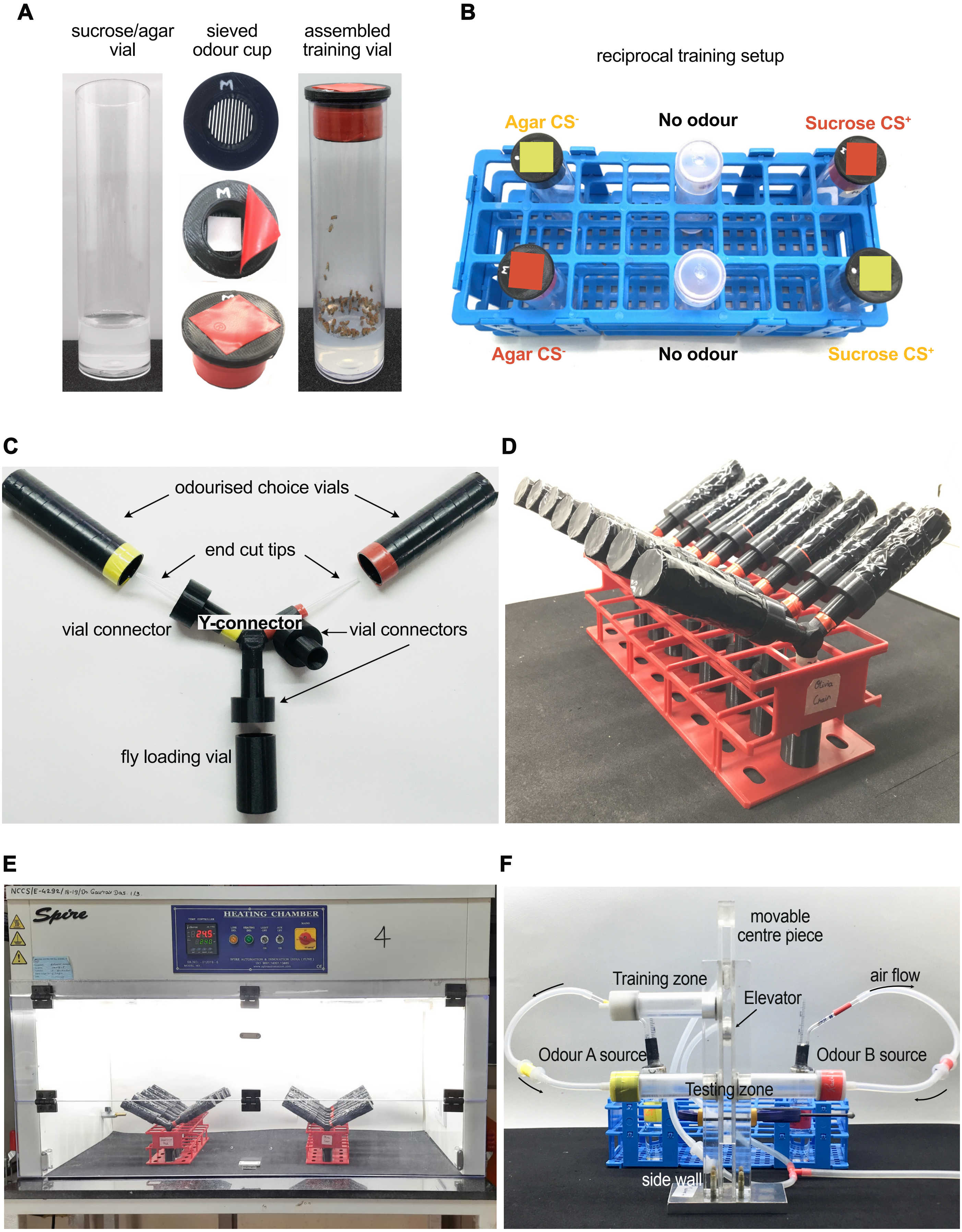

**Supplementary Figure 1** ***(accompanying Figure 1)*****: Parts and assembly of training and testing setups. (A)** Training vial with odour cup used for conditioning. **(B)** Arrangement of training vials in a plastic rack during training. **(C)** Assembly of the Y-maze from standard fly plasticware and 3D printed parts. **(D)** Vertical arrangement of Y-mazes in a plastic rack during testing. **(E)** Custom designed temperature-controlled chamber used for behaviour assays. **(F)** A typical T-maze.

**Supplementary Figure 2**

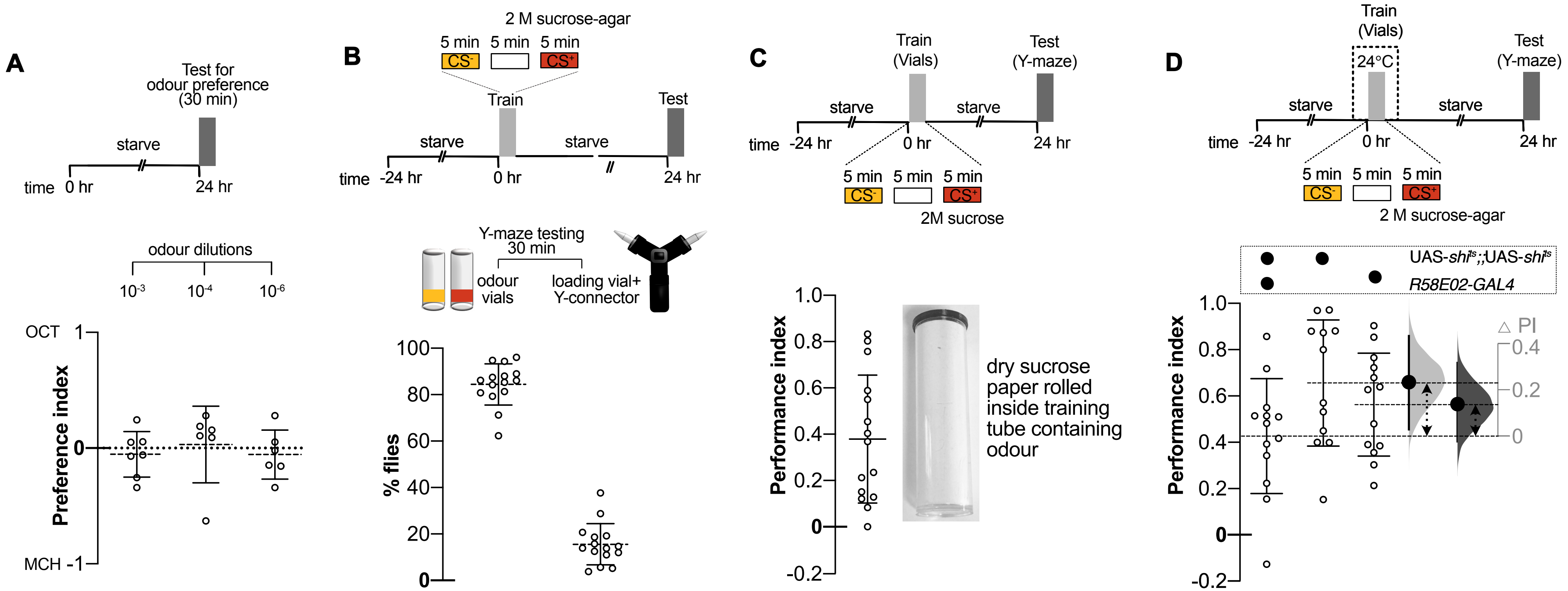

**Supplementary Figure 2** ***(accompanying Figure 1 and 2)*****: Training and testing in the Y-mazes.** **(A)** Groups of starved and naive wild type CS flies show equal distribution amongst the two odour choice vials at ~$10^{-3}$, ~$10^{-4}$ and ~$10^{-6}$ fold dilutions of both OCT and MCH (*n=6* for all groups). **(B)** The majority of flies (~85 %) participate in olfactory choice after testing for 30 min. The number of flies in the choice vials versus the rest of the assembly is quantified *(n=15)*. **(C)** Significant 24 h memory performance is seen after training flies with 2.66 M sucrose solution dried on filter paper (*n=15; $CS^+$ OCT=8, MCH*=7). The sucrose paper lined the training vials. **(D)** Room temperature control experiment for (Figure 2C) shows that the experimental genotype, *R58E02-GAL4/UAS-$Shibire^{ts1}$ (n=13; $CS^+$ OCT=7; $CS^+$ MCH=6)* performs as well as the parental controls *UAS-$Shibire^{ts1}$(n=12;$CS^+$ OCT=7, $CS^+$ MCH=5)* and *R58E02-GAL4 (n=13;$CS^+$ OCT=7, $CS^+$ MCH=6)*. Effect size or the difference in mean performance index (ΔPI) plots are shown for all. They show the observed effect size of the experimental group compared to a control (large black dots), 95 % CI (end of vertical error bars) and the bootstrap resampling error (curve) of effect size. Double-headed dashed arrows mark the groups being compared

#### Supplementary Figure 3

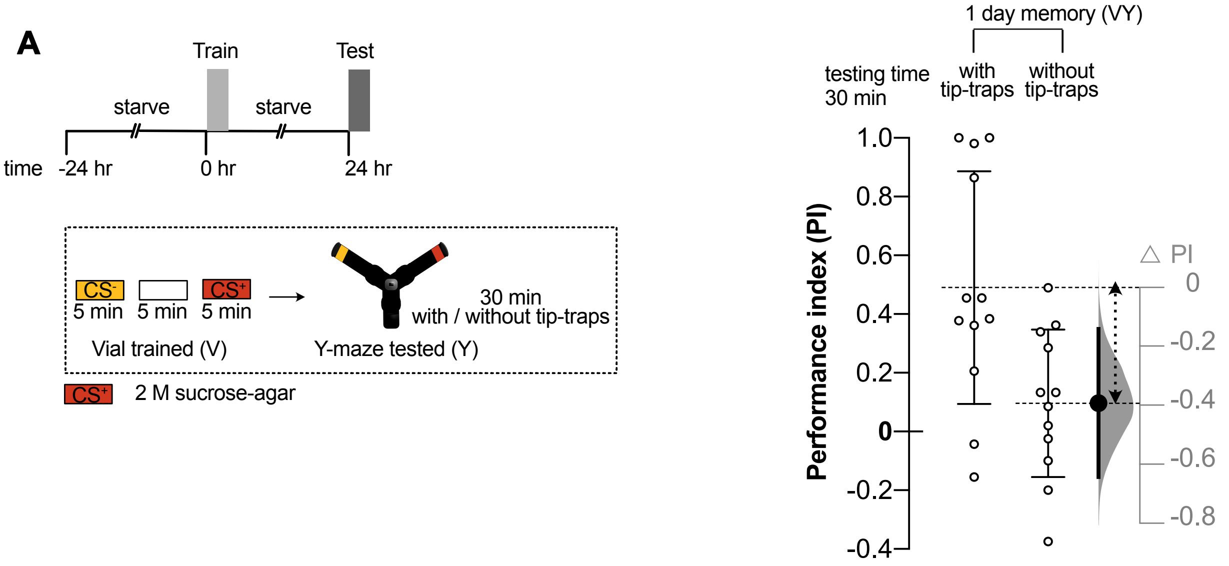


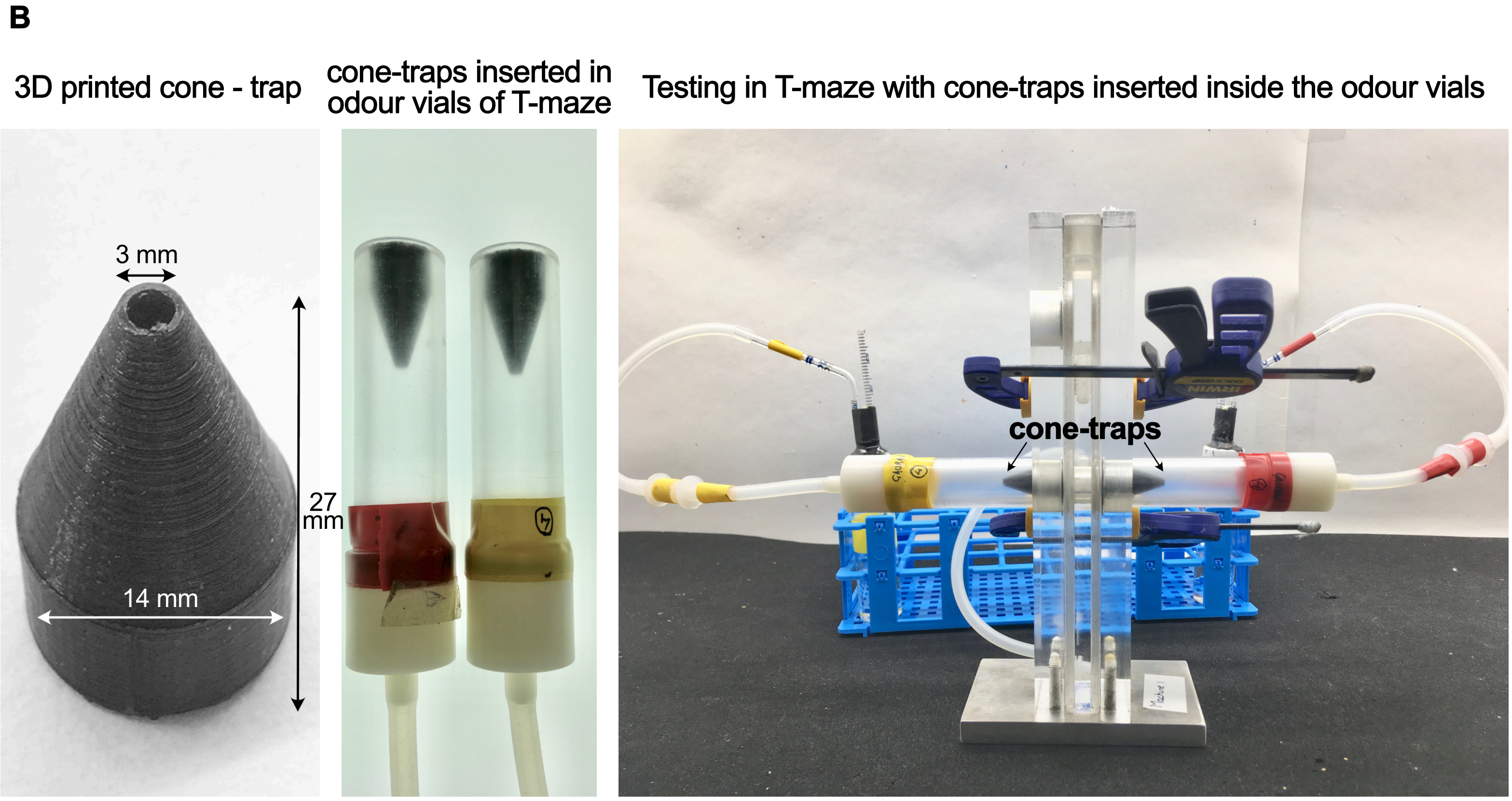

**Supplementary Figure 3 (*accompanying Figure 3)* 1-day memory retrieval is improved when flies are restricted to their first committed choice using traps**

**(A)** 1-day memory performance of flies trained with 2 M sucrose in vials and tested in Y-maze with or without tip-traps for 30 min. Compared to flies tested with tip-traps in Y-maze (*n=12; $CS^+$ OCT=6, MCH*=6.), memory disruption observed in flies tested without tip-traps in Y-maze (*n=12; $CS^+$ OCT=6, MCH*=6).Effect size or the difference in mean performance index (ΔPI) plots are shown for all. They show the observed effect size of the experimental group compared to a control (large black dots), 95 % CI (end of vertical error bars) and the bootstrap resampling error (curve) of effect size. Double-headed dashed arrows mark the compared groups. **(B)** 3D printed conical traps fixed at the open end of the odour vials used for choice in a T-maze.

### Supplementary Figure 4

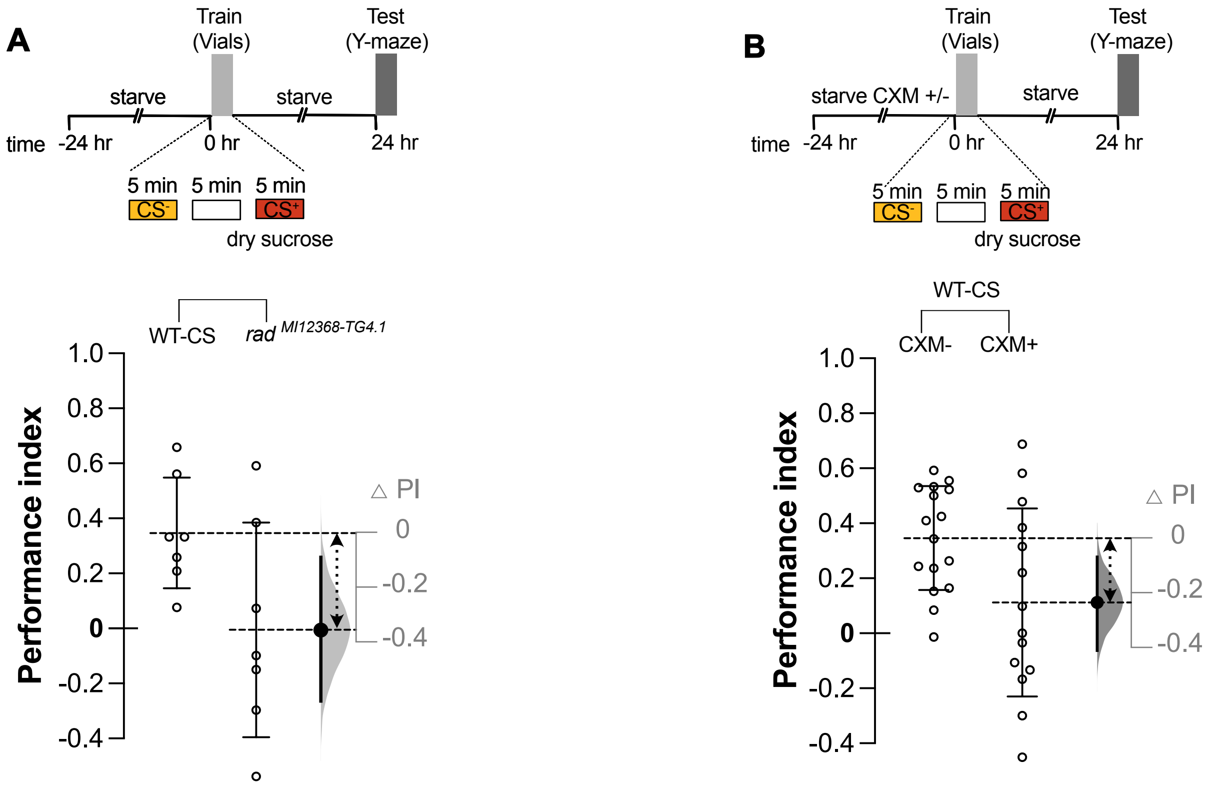

**Supplementary 4 *(accompanying Figure 4)*: Susceptibility of 24 h dry sucrose memory upon *rad*$^{MI12368\text{-}TG4.1}$ mutation and CXM treatment (A)** 1 day dry sucrose memory disrupted in *rad*$^{MI12368\text{-}TG4.1}$ mutants *(n=7; CS$^+$ OCT=4, MCH=3)* when compared to WT-CS flies *(n=;7 CS$^+$ OCT=4, MCH=3)*. Flies were trained in vials and tested in Y-maze. **(B)** 1 day memory comparison upon CXM treatment in flies trained in vials with dry sucrose and tested in Y-maze. Memory is disrupted in WT-CS CXM$^+$ groups *(n=14; CS$^+$ OCT=7, MCH=7)* compared to WT-CS CXM$^-$ groups *(n=16; CS$^+$ OCT=8, MCH=8)*. To the right of each graph, effect size or the difference in mean performance index (ΔPI) plots are shown. These plots depict the bootstrap sampling error distribution curve of ΔPI. The actual mean difference of the groups is depicted as a large black dot and the 95 % CI is indicated by the ends of the vertical error bars. Double-headed dashed arrows between the group-means indicate the groups being compared.

**Supplementary Table 1 (ST1): Details of statistical analysis performed in GraphPad Prism version 8.3.1.**

| Figure | Experiment name | Statistical test (GraphPad Prism) | *p-value* | Estimation plot used https://www.estimationstats.com/#/ |
|---|---|---|---|---|
| 1B | **Varying odour dilutions used for training and testing** | | | |
| | $10^{-3}$ odour dilution vs $10^{-4}$ odour dilution | Tukey's multiple comparisons | 0.0023 | Shared control |
| | $10^{-3}$ odour dilution vs $10^{-6}$ odour dilution | | <0.001 | |
| | $10^{-4}$ odour dilution vs $10^{-6}$ odour dilution | | 0.0131 | |
| 1C | **Varying odour-reward association time for training** | | | |
| | 5 min vs 2 min | Unpaired t-test with Welch's correction | 0.8364 | Two-groups |
| 1D | **Varying sucrose concentration for training** | | | |
| | 2 M vs 100 mM | Tukey's multiple comparisons | 0.0512 | Shared control |
| | 2 M vs 50 mM | | 0.0109 | |
| | 100 mM vs 50 mM | | 0.7569 | |
| 1E | **Testing memory performance at different time points** | | | |

| | | | | |
|---|---|---|---|---|
| | 1 day vs 3 day | Tukey’s multiple comparisons | 0.3001 | Shared control |
| | 1 day vs 5 day | | 0.0833 | |
| | 1 day vs 7 day | | 0.2521 | |
| | 3 day vs 5 day | | 0.9019 | |
| | 3 day vs 7 day | | 0.9822 | |
| | 5 day vs 7 day | | 0.9957 | |
| 2A | **24 h memory performance in memory mutant flies** | | | |
| | WT-CS vs *dunce* | Dunnet’s multiple comparisons | 0.0007 | Shared control |
| | WT-CS vs *tequila* | | 0.0475 | |
| | WT-CS vs $rad^{MI12368\text{-}TG4.1}$ | | 0.0131 | |
| 2B | **24 h memory performance in *dumb* flies** | | | |
| | WT-CS vs $dumb^{1}$ | Dunnet’s multiple comparisons | 0.002 | Shared control |
| | WT-CS vs $dumb^{2}$ | | <0.001 | |
| 2C | ***R58E02* PAM dopaminergic neurons silencing at 32℃** | | | |
| | *UAS-Shi$^{ts1}$ vs R58E02-GAL4* | Tukey’s multiple comparisons | 0.6822 | Shared control |
| | *UAS-Shi$^{ts1}$-R58E02-GAL4* vs *UAS-Shi$^{ts1}$* | | <0.001 | |
| | *UAS-Shi$^{ts1}$-R58E02-GAL4* vs *R58E02-GAL4* | | <0.001 | |

| | | | | |
|---|---|---|---|---|
| Supp 2D | ***R58E02* PAM dopaminergic neurons silencing at 24℃** | | | |
| | *UAS-Shi$^{ts1}$ vs R58E02-GAL4* | Tukey's multiple comparisons | 0.6176 | Shared control |
| | *UAS-Shi$^{ts1}$-R58E02-GAL4* vs *UAS-Shi$^{ts1}$* | | 0.0670 | |
| | *UAS-Shi$^{ts1}$-R58E02-GAL4* vs *R58E02-GAL4* | | 0.3521 | |
| | *UAS-Shi$^{ts1}$;;UAS-Shi$^{ts1}$* (PI= -0.4615) | Outlier test ROUT (Q=10%) | | |
| 3A | **Comparison of training and testing in T-maze and Y-maze** | | | |
| | VY vs VT | Tukey's multiple comparisons | 0.0586 | Two groups |
| | VY vs TY | | 0.9342 | |
| | VY vs TT | | 0.0310 | |
| | VT vs TY | | 0.1460 | |
| | VT vs TT | | 0.9906 | |
| | TY vs TT | | 0.0801 | |
| 3B | **Comparison of 5-day memory in T-maze and Y-maze** | | | |
| | 1 day: T-maze vs Y-maze | Unpaired T-test with welch's correction | 0.0052 | Two groups |
| | 5th day: Tmaze vs Y-maze | | 0.0006 | |
| | 5th day Y-maze testing (PI= -0.447) | Outlier test ROUT (Q=10 %) | | |

| | | | | |
|---|---|---|---|---|
| 3C | **Comparing between 5 min and 30 min testing time in Y-maze on 1 day and 5 day memory** | | | |
| | 1 day: 30 min vs 5 min | Unpaired T-test with welch's correction | 0.2583 | Two groups |
| | 5th day: 30 min vs 5 min | | 0.3996 | |
| | 1 day: 30 min vs 5 min<br>1 day 30 min - (PI= -0.014) and (PI= 0.2783) | Outlier test<br>ROUT (Q=10 %) | | |
| 3D | **Comparison of 1 day memory, tested in Y-maze for 5 min with or without tip-traps** | | | |
| | With tip-traps vs without tip-traps (5 min, Y-maze) | Unpaired T-test with welch's correction | 0.0630 | |
| | With tip-traps vs without tip-traps (5 min, Y-maze)<br>With tip-traps:(PI= -1) | Outlier test<br>ROUT (Q=10 % | | |
| 3E | **Comparison of 1 day memory, tested in T-maze with or without tip-traps** | | | |
| | With tip-traps vs without tip-traps (5 min) | Tukey's multiple comparisons | 0.0439 | |
| | With tip-traps (5 min) vs without tip-traps (2 min) | | 0.3957 | |
| | Without tip-traps (5 min) vs without tip-traps (2 min) | | 0.8590 | |
| Supp | **Comparison of 1 day memory, tested in Y-maze for 30 min with or** | | | |

| | | | | |
|---|---|---|---|---|
| 3A | **without tip-traps** | | | |
| | With tip-traps vs without tip-traps (30 min, Y-maze) | Unpaired T-test with welch's correction | 0.009 | |
| 4A | **Comparing aversive memory scores in Y-mazes versus T-mazes** | | | |
| | D-Arabinose vs $CuSO_4$ +D-Arabinose (Y-maze) | Unpaired t-test with Welch's correction | <0.001 | Two groups |
| | D-Arabinose vs $CuSO_4$ + D-Arabinose (T-maze) | | 0.0966 | |
| 4B | **Protein synthesis dependence of sweet and aversive taste memories** | | | |
| | D-Arabinose $CXM^-$ vs D-Arabinose $CXM^+$ | Tukey's multiple comparison | 0.2300 | Two groups |
| | D-Arabinose $CXM^-$ vs $CuSO_4$ + D-Arabinose $CXM^-$ | | 0.0018 | |
| | D-Arabinose $CXM^-$ vs D-Arabinose + $CuSO_4$ $CXM^+$ | | 0.3579 | |
| | D-Arabinose $CXM^+$ vs D-Arabinose + $CuSO_4$ $CXM^-$ | | 0.1959 | |
| | D-Arabinose $CXM^+$ vs D-Arabinose + $CuSO_4$ $CXM^+$ | | 0.9986 | |

| | | | | |
|---|---|---|---|---|
| | D-Arabinose + $CuSO_4$ CXM$^-$ vs D-Arabinose + $CuSO_4$ CXM$^+$ | | 0.1788 | |
| 4C | **Protein synthesis dependent memory in WT-CS and *rad*$^{MI12368-TG4.1}$ mutants** | | | |
| | *rad*$^{MI12368-TG4.1}$ CXM$^-$ vs *rad*$^{MI12368-TG4.1}$ CXM$^+$ | Unpaired T-test with welch's correction | 0.210 | Two groups |
| | *WT-CS* CXM$^-$ vs *WT-CS* CXM$^+$ | Unpaired T-test with welch's correction | 0.2775 | |
| | *WT-CS* CXM$^-$ vs *WT-CS* CXM$^+$<br>*WT-CS* CXM$^+$ (PI= -0.697) | Outlier test ROUT (Q=10 % | | |
| Supp 4A | **1 day dry sucrose memory in *rad*$^{MI12368-TG4.1}$ mutants when tested in Y-maze** | | | |
| | WT-CS vs *rad*$^{MI12368-TG4.1}$ | Unpaired T-test with welch's correction | 0.0628 | Two groups |
| Supp 4B | **Susceptibility of 1 day dry sucrose memory upon CXM treatment when tested in Y-maze** | | | |
| | *WT-CS* CXM$^-$ vs *WT-CS* CXM$^+$ | Unpaired T-test with welch's correction | 0.034 | Two groups |
